# An amphipathic helix enables septins to sense micron-scale membrane curvature

**DOI:** 10.1101/379982

**Authors:** Kevin S. Cannon, Benjamin L. Woods, John M. Crutchley, Amy S. Gladfelter

## Abstract

The geometry of cells is well described by membrane curvature. Septins are filament forming, GTP-binding proteins that assemble on positive, micrometer-scale curvatures. Here, we examine the molecular basis of curvature sensing by septins. We show that differences in affinity and the number of binding sites drive curvature-specific adsorption of septins. Moreover, we find septin assembly onto curved membranes is cooperative and show that geometry influences higher-order arrangement of septin filaments. Although septins must form polymers to stay associated with membranes, septin filaments do not have to span micrometers in length to sense curvature, as we find that single septin complexes have curvature-dependent association rates. We trace this ability to an amphipathic helix (AH) located on the C-terminus of Cdc12. The AH domain is necessary and sufficient for curvature sensing both *in vitro* and *in vivo*. These data show that curvature sensing by septins operates at much smaller length scales than the micrometer curvatures being detected.

## Introduction

Shape is a fundamental feature in cell biology and can be thought in terms of membrane curvature (Zimmerberg and Kozlov, 2006; Cannon, Woods and Gladfelter, 2017). Cellular membrane curvature is a continuum, spanning nanometer to micrometer scales. How do cells use nanometer-sized components to perceive micron-scale changes in shape? Septins, are filament-forming, GTP-binding proteins that localize to sites of micron-scale membrane curvature from yeast to humans (Field *et al*., 1996; Pan, Malmberg and Momany, 2007; Bridges *et al*., 2016). Examples of curvature-associated localizations include the bud-neck in *S. cerevisiae* (Byers and Goetsch, 1976; Haarer and Pringle, 1987; Ford and Pringle, 1991), bases of dendritic spines in neurons (Cho *et al*., 2011), branches in filamentous fungi (Westfall and Momany, 2002; DeMay *et al*., 2009; Bridges *et al*., 2016), and the cytokinetic furrow (Spiliotis, Kinoshita and Nelson, 2005; Joo, Surka and Trimble, 2007; Maddox *et al*., 2007). At these sites, septins coordinate cell cycle progression (Longtine *et al*., 2000; Sakchaisri *et al*., 2004), influence diffusion in the membrane (Clay *et al*., 2014; Yamada *et al*., 2016), and act as a scaffold to recruit proteins required for chromosome segregation (Spiliotis, Kinoshita and Nelson, 2005) and cytokinesis (Meitinger *et al*., 2011; Finnigan *et al*., 2015).

Septins carry out these functions by assembling into heteromeric, rod-shaped, non-polar complexes which can anneal end-on to polymerize into filaments at the plasma membrane (Field *et al*., 1996; John *et al*., 2007; Sirajuddin *et al*., 2007; Bertin *et al*., 2008). Budding yeast possess five mitotic septins that assemble into hetero-octamers in which the terminal subunit is either Cdc11 or Shs1(Garcia *et al*., 2011; Khan *et al*., 2018). Purified recombinant septins from yeast and humans preferentially adsorb onto micrometer curvatures in the absence of any cellular factors (Bridges *et al*., 2016) indicating that curvature sensing is a conserved feature of the septin cytoskeleton. The mechanism underlying how septins sense micron-scale membrane curvature is unclear.

Most of what we know about curvature sensing comes from nanometer-sized molecules interacting with nanometer-scale curvatures. Proteins containing BAR domains and/or amphipathic helices (AH) utilize combinations of membrane insertion (Drin and Antonny, 2010), oligomerization, and scaffolding mechanisms (Simunovic, Srivastava and Voth, 2013; Simunovic *et al*., 2015) to either sense or deform the local curvature. Membrane curvature generates lipid packing defects, providing binding sites for AHs (Hatzakis *et al*., 2009). Proteins such as α-synuclein (Pranke *et al*., 2011), Opi1 (Hofbauer *et al*., 2018), and ArfGAP1 (Drin *et al*., 2007) employ this mechanism to sense curvature. Interestingly, known micrometer-scale curvature sensors including SpoVM (Ramamurthi *et al*., 2009), MreB (Ursell *et al*., 2014; Hussain *et al*., 2018) contain AHs.

In this study, we investigate the mechanisms of septin curvature sensing. We discovered that septins vary in their affinity for different curvatures, that single septin oligomers bind with different association rates depending on the curvature and that a conserved AH is both necessary and sufficient for curvature sensing by septins. This study provides the first insights into the molecular basis for how septins sense curvature.

## Results and Discussion

### Analysis of septin saturation binding to different curvatures

We first generated saturation binding isotherms from which affinity, maximal binding and cooperativity can be estimated for septins on different curvatures. We used a minimal reconstitution system consisting of recombinant yeast septin complexes (Cdc11-GFP, Cdc12, Cdc3, and Cdc10) and supported lipid bilayers (SLBs) formed on silica beads of different curvatures (Gopalakrishnan *et al*., 2009; Bridges *et al*., 2016). We measured septin binding onto SLB-coated beads at a range of concentrations with different bead sizes using quantitative microscopy. We found that septins have the strongest affinity for 1 μm beads (curvature, κ= 2 μm^−1^, K_d_ 13.5 nM), followed by 3 μm beads (κ= 0.67 μm^−1^, K_d_ 18.5 nM), and 0.5 μm beads (κ= 4 μm^−1^, K_d_ 34.3 nM) (Fig.1, Table 1). Additionally, the difference in maximal binding capacity (B_max_) is dramatically different for tested bead sizes, indicating that there are curvature-dependent differences in the number of effective binding sites for septins (Fig.1, Table 1). This is not due to differences in surface area on different bead sizes, as we normalized for surface area. At high septin concentrations, filaments formed in solution at curvature κ= 4 μm^−1^, indicating excess complex is available for polymerization, suggesting the number of binding sites is limiting at this curvature (Fig.1). Hill coefficients of 2.6 (κ= 4 μm^−1^ and κ= 0.67 μm^−1^) and 2.9 (κ= 2 μm^−1^), indicated septin adsorption on all tested membrane curvatures is cooperative, consistent with the observation that some beads are fully bound by septins whereas others have none.

**Figure 1.**
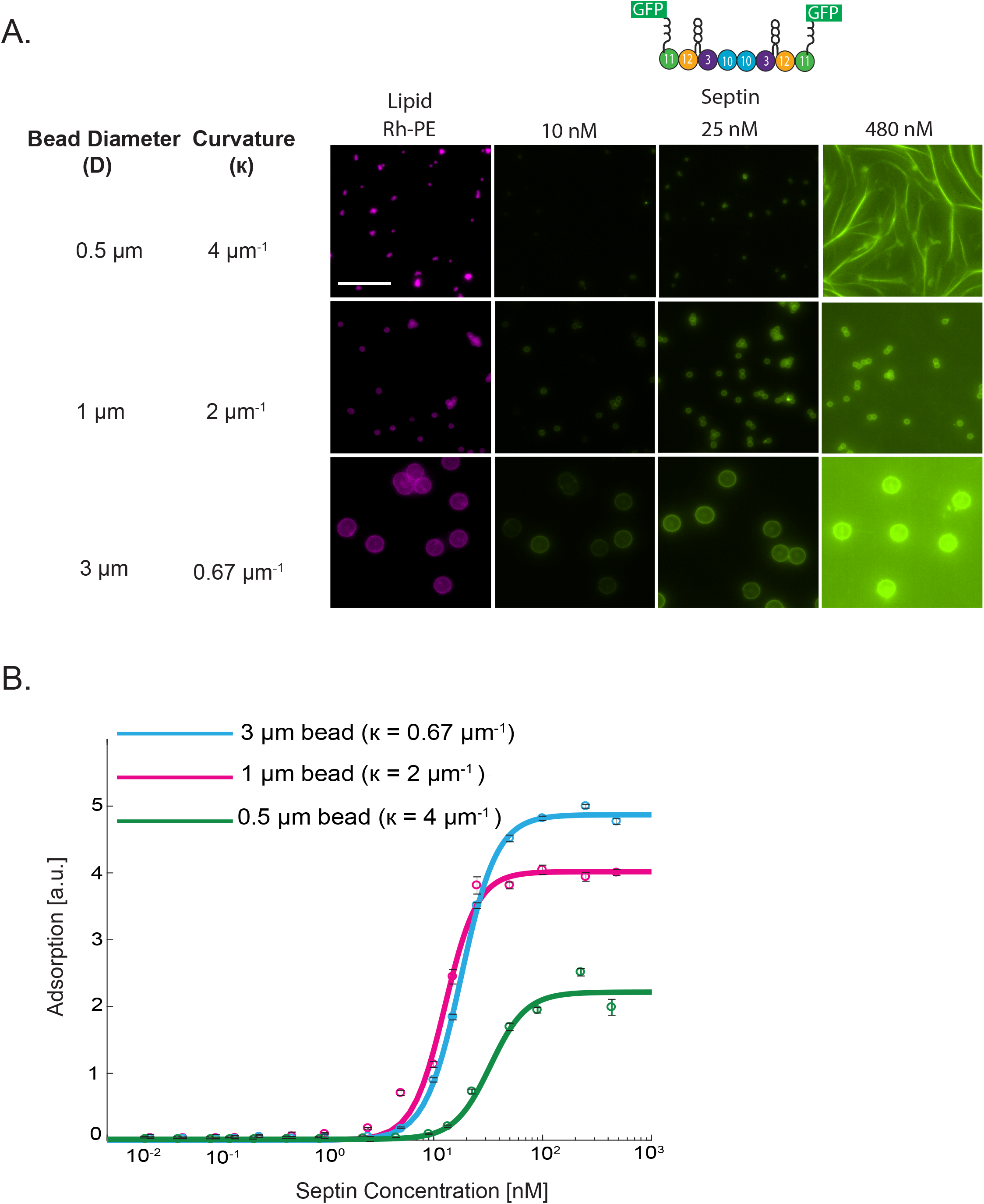
Septins bind cooperatively to curved membranes with differences in affinity and maximal binding. SLBs (75% DOPC, 25% (PI), and trace amounts of Rh-PE) were reconstituted on silica beads. Purified septins were added at several concentrations through saturation. (A) Representative images are maximum intensity projections. Scale bar 10 μm. (B) Quantification of septin adsorption at equilibrium onto SLBs of varying curvature. Each data point represents the mean intensity for 98-600 beads. Error bars are SEM, N=3.

**Table 1.**
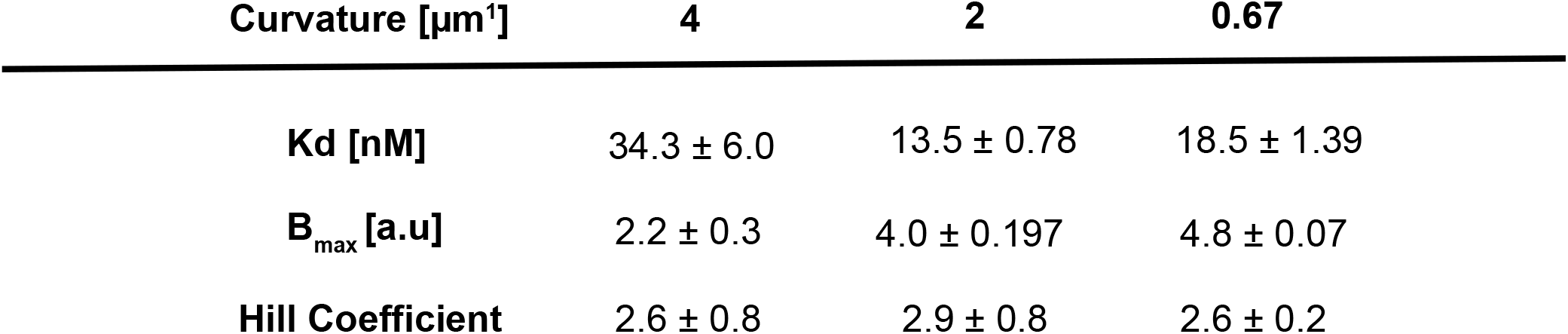
Biochemical parameters for septin adsorption onto various membrane curvatures

We predict that affinity is a major driver of curvature sensing at low septin concentrations and at high concentrations how dense filaments can arrange on different curvatures becomes important. Collectively, these data suggest that differences in septin affinity and the number of available binding sites can drive preferential assembly of septins onto specific curvatures in a cooperative fashion. Cooperativity in assembly could arise at filament formation, lateral associations between filaments and/or other higher-order structures including layers of filaments.

### Septin filament orientation is dependent on membrane curvature

To evaluate how cooperativity may arise at the level of polymerization, we asked how long a septin filament must be to align along optimum curvature. We adapted the SLB assay to rods of different diameters to visualize filaments using scanning electron microscopy (SEM). We scored the length and orientation of filaments on a range of rod diameters binned to three categories (100-400 nm, 401-600 nm, and 601-1000 nm). For all three categories, we found that single septin octamers (32 nm in length) sampled a variety of orientations as each image is a snapshot at the time of fixation (Fig.2E-G, orange boxes, Table 2). For rod diameters of 600-1000 nm, filaments 64 nm long tend to align along the axis of principal curvature (Fig.2G, green boxes). On rod diameters from 401-600 nm, septin filaments up to 128 nm long (four annealed octamers) had a wide distribution of orientations (Fig.2F). In contrast, for the smallest rods with diameters spanning 100-400 nm, 64 nm filaments begin to align along the long-axis (axis of zero curvature) of the rod to avoid the higher positive curvature (Fig.2E). For longer filaments, this alignment is even more evident (Fig.2E and G, pink and blue boxes). These data suggest even short septin filaments align along positive curvature.

**Figure 2.**
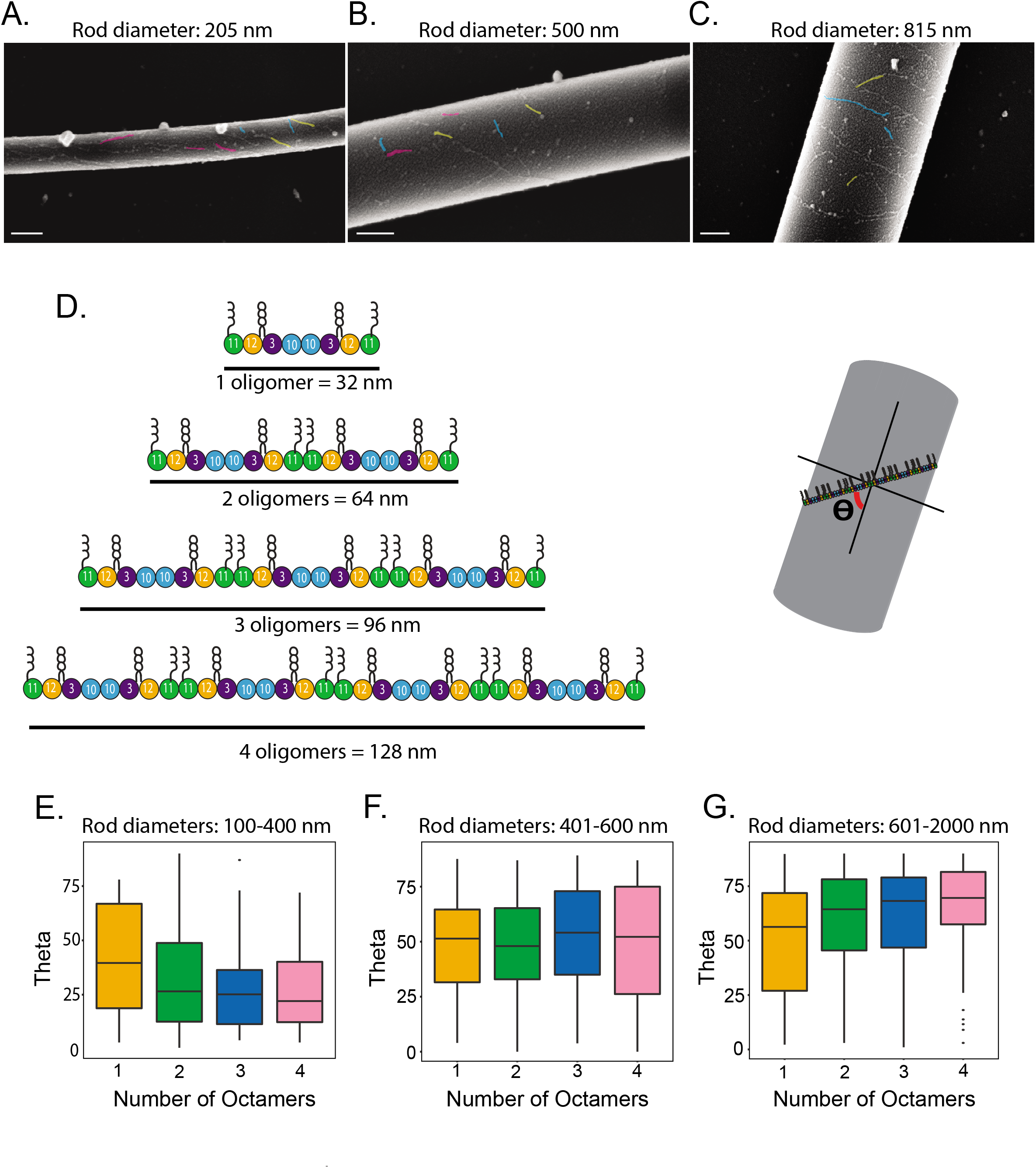
Septin filament alignment towards the axis of principal curvature is dependent on filament length. SLBs (75% DOPC, 25% PI, and trace amounts of Rh-PE) were reconstituted on borosilicate rods of different diameters ranging from approximately 100 nm to 2300 nm.(A-C) Representative images of septin filament alignment on rods from the three different categories. Scale bar 200 nm. A subset of filaments was false colored to depict alignments; pink are parallel to the long axis of curvature; yellow are oriented at ~ 45° blue are aligned to the axis of principal curvature. (D) Schematic of septin filament length in terms of the number of octamers and a cartoon depicting how filament orientation relative rod was measured. (E-G) Box and whisker plot quantifying septin filament alignment on various rod diameters as a function of filament length binned to three diameter ranges. Black bars represent the median. Error bars are standard deviation (E) 100-400 nm rods. N= 23 rods and 193 filaments (F) 401-600 nm rods. N= 15 rods and 189 filaments. (G) 601-2000 nm rods. N= 24 rods and 491 filaments.

**Table 2.**
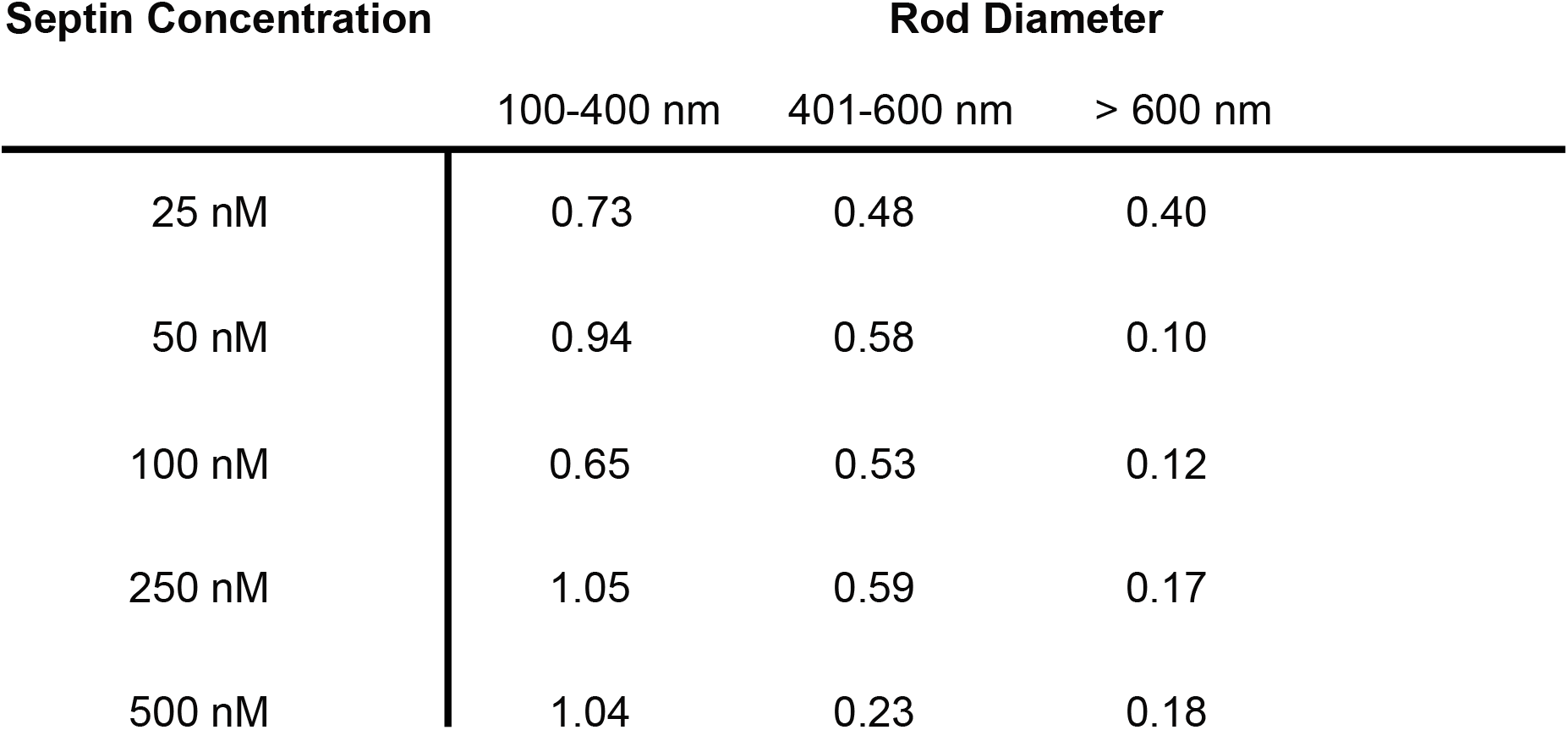
Coefficient of variation for septin filament alignment on rods

We predicted that with longer septin filaments and/or lateral interactions that a tighter distribution of orientations would be seen. Indeed, on narrow and wide diameter rods (Fig.3E, pink and blue) with septins at high concentrations (50-500 nM) filaments are well aligned with one another and tightly packed. On rod diameters from ~400 nm through ~ 630 nm, septin filaments exhibit a wide range of orientations (Fig.3E, yellow). However, neighboring filaments have a tendency to align with one another. Septin filament orientation is very similar on other lipid compositions including DOPS, DOPE, PI (4,5)P_2_, showing that septins can sense curved membranes on a variety of lipid compositions (Supplemental Fig.2). Thus, at high density, septin filaments more closely match the curvature of the rod than septin filaments at a low-density.

**Figure 3.**
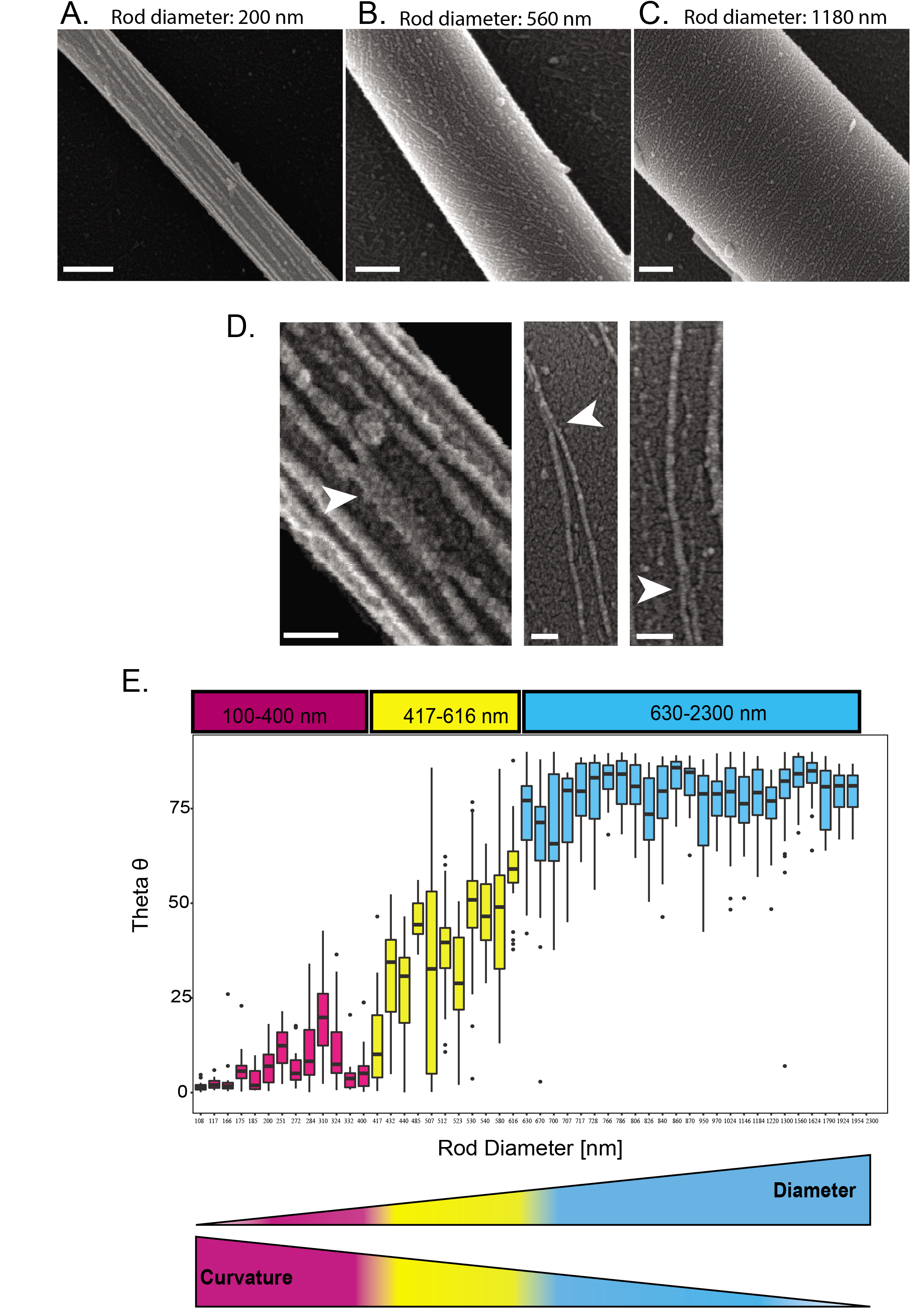
Septin filament orientation is dependent on membrane curvature. SLBs (75% DOPC, 25% PI, and trace Rh-PE) were generated on borosilicate rods of different diameters (100-2300 nm). Purified septins were added to SLBs at saturating concentrations (from 50 nM to 500 nM) and imaged using SEM. (A-C) Representative images of septin filament alignment on rods from the three different categories at 50 nM septin concentration. (D) Example septin bundling (white arrows). (E) Box and whisker plots of septin filament orientation on measured rod diameters at several septin concentrations (50 nM, 100 nM, 250 nM, and 500 nM). The number of filament orientations measured per rod ranged > 10 filaments. Quantification is from 49 rods. Measured filaments were greater than 100 nm (~3 octamers) in length. Black bars represent the median. Error bars are standard deviations.

These data suggest that the ability of septins to perceive micron-scale curvature may be in part driven by septin filament density and filament length. Both length and packing may contribute to the cooperativity detected in the saturation binding curves (Fig.1B). Aligned septin filaments in close proximity to each other might be stabilized by lateral interactions with one another as has been noted previously (Frazier *et al*., 1998; Bertin *et al*., 2008; Sadian *et al*., 2013) (Fig.3D). We also imagine that tightly packed septins might sterically hinder one another, thus “enforcing” alignment along the optimal curvature through transient interactions while diffusing on the membrane. At higher septin concentrations, filaments are also longer on rods than at low concentrations so it is possible that filament length is also a driving factor in this assay.

### Association rates of single septin complex are curvature sensitive

We next examined curvature-dependent septin affinity differences. We measured the number and duration of binding events to calculate association and off rates for single septin complexes on curvatures of 2 μm^−1^ and 0.67 μm^−1^ (Fig.4, Supplemental Fig. 1). Histograms of dwell times for single septin-complex binding events were fit to a single exponential decay function. The measured dwell times for these curvatures were 0.21s ± 0.019 and 0.21 ± 0.023, for κ = 2 μm^−1^ and κ =0.67 μm^−1^, respectively (N_particles per curvature_ > 150). Dwell times correspond to a high off-rate, supporting previous observations that a single septin octamer does not stably associate with membranes (Bridges *et al*., 2016). Notably, the measured dwell times were the same on both curvatures tested, suggesting that curvature does not influence the off rate. Next, association rates were quantified by calculating the number of binding events over the product of septin concentration, time, and binding area. We found that single septin octamers show differences in the association rate as a function of membrane curvature (Fig.4, κ = 2 μm^−1^ was 0.061 μm^−2^s^−1^nM^−1^ and κ = 0.67 μm^−1^ was 0.025 um^−2^s^−1^nM^−1^).

**Figure 4.**
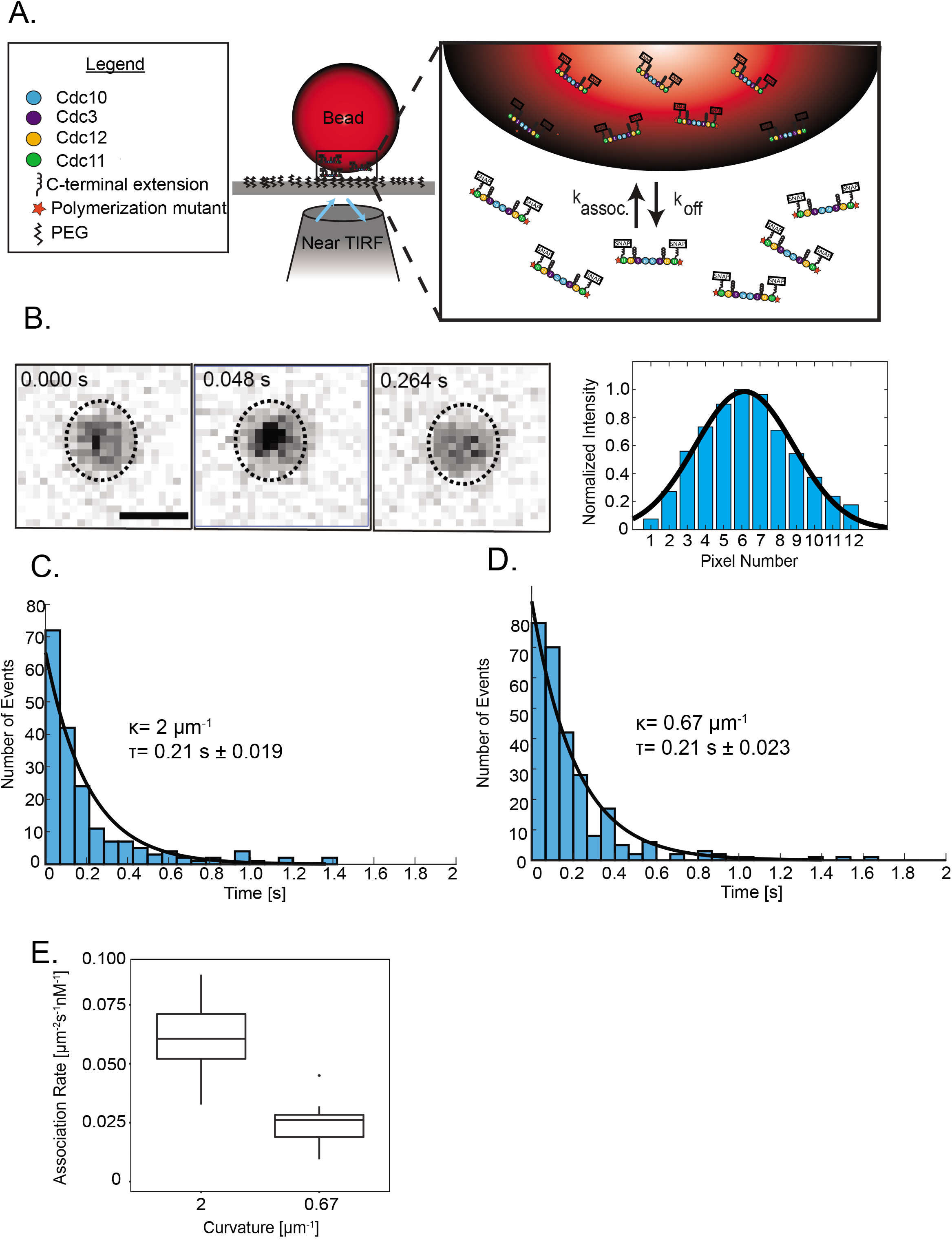
Single septin complexes have a higher association rate for optimal membrane curvatures. (A) SLBs (75% DOPC and 25% PI) were reconstituted on curvatures of 2 μm^−1^ or 0.67 μm^−1^ and flowed between two polyethylene-glycol (PEG, black) coated coverslips (top coverslip not shown). Non-polymerizable septin complexes were then flowed into the chamber and septin binding and unbinding events were observed using near-TIRF microscopy. (B) Representative images of a binding event on κ= 2 μm^−1^ (1 μm bead). Line scan through the particle shows signal intensity fit to a Gaussian function. (C-D) Dwell time histograms for membrane curvature of 2 μm^−1^ and 0.67μm^−1^, respectively. (E) Box and whisker plot quantifying association rate of non-polymerizable septin complexes onto both curvatures (κ =2μm^−1^ and 0.67μm^−1^). Black bars represent the median. Error bars are standard deviations. N_beads_ = 10 (for κ =2μm^−1^ and 0.67μm^−1^) N_binding events_ = 844 and 315 for for κ =2μm^−1^ and 0.67μm^−1^, respectively.

The ~2.5-fold difference in association rate between these two curvatures was striking, given that our saturation binding isotherms show K_d_ values so close to one another (13.5 and 18.5 nM, respectively). This could be explained by the fact that our binding curve data is a summation of several reactions: initial binding, septin annealing and fragmentation events, and lateral interactions, all of which may be influenced by curvature and contribute to the K_d_ values. These results suggest that even a single septin octamer can detect curvature differences through the initial association with the curved membrane and curvature-dependent bilayer defects are perceptible on micron-scale curvature.

### An amphipathic helix domain in septins is necessary and sufficient for curvature sensing

How can the association rate of a single septin octamer be sensitive to curvature on the micron scale? Many proteins that sense nanometer scale curvatures contain membrane-binding amphipathic helices (Drin *et al*., 2007; Drin and Antonny, 2010). Interestingly, another micron scale curvature sensor, SpoVM, a 26-amino acid polypeptide, utilizes an 11-residue amphipathic helix to sense curvature (Gill *et al*., 2015) and also shows curvature-dependent association rates (Kim *et al*., 2017). We manually searched for AH motifs in septin sequences using Heliquest (Gautier *et al*., 2008) to screen primary sequences for α-helical properties and found conserved, predicted amphipathic helices at the C-termini of a subset of septins (Fig.5). Despite differences in amino acid composition, septin AH motifs are similarly situated within the primary sequence and have similar lengths, net charges, and hydrophobic moments, suggesting that the physicochemical properties of the AH domains are highly conserved across multiple species (Supplemental Fig.3).

**Figure 5.**
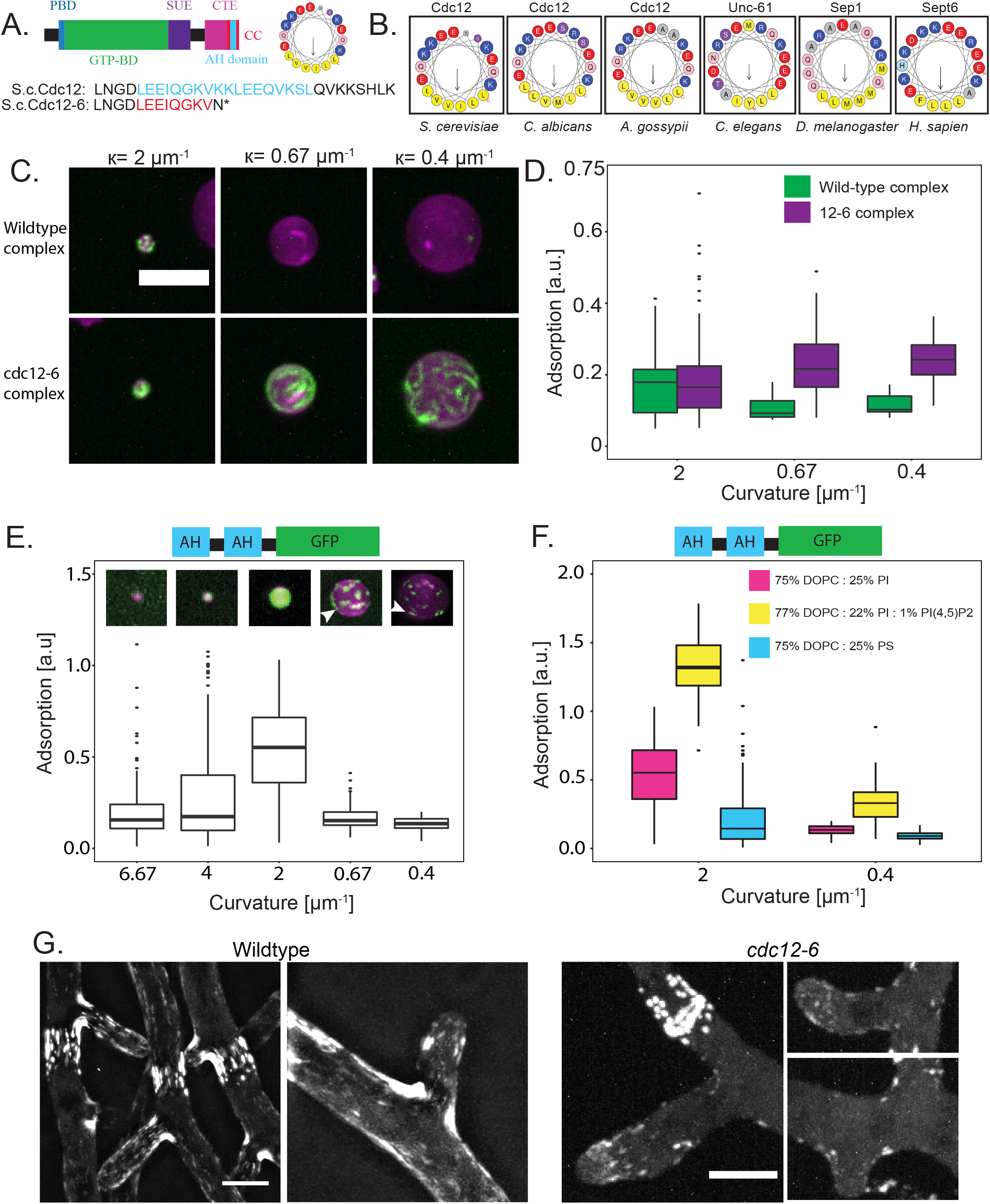
Septins have conserved amphipathic helices at their C-termini. (A) Domains within the yeast septin, Cdc12. PBD (blue): polybasic domain; GTP-BD (green): GTP-binding domain; SUE (purple): Septin unique element; CTE(pink): C-terminal extension; CTE(magenta): coiled-coil domain (red); AH domain (cyan). The AH domain is shown as a helical wheel corresponding primary sequences for wild-type Cdc12 and Cdc12-6 (below). (B) Amphipathic helices found in the C-terminal extensions of septins in multiple species represented through helical wheel diagrams. (C) 1 nM of purified wild-type septin complexes (panel 1) and *cdc12-6* septin complexes (panel 2) adsorption (green) onto SLBs (magenta) with different curvatures. (D) Box and whisker plot quantifying adsorption of wild-type or cdc12-6 septin complexes onto SLBs of various curvature. Black bars represent the median. Error bars represent the standard deviation. n > 50 beads for each curvature. (E) Representative images of 1 μM of 2x-Cdc12 AH-GFP binding onto SLBs on beads with curvatures. Arrows highlight filament-like structures. (F) Box and whisker plot of 2x-Cdc12 AH-GFP adsorption onto different membrane compositions and curvatures. Black bars represent the median. Error bars at each curvature are standard deviations for n> 100 beads. (G) Septins assemble at the base hyphal branches in wild-type cells (wildtype; Cdc11a-GFP), but rarely assemble in *cdc12-6* mutants (cdc12-6-GFP) at permissive temperature.

We asked if the septin AH domain confers specificity for curved membranes. Interestingly, the AH domain of Cdc12 is truncated in a well characterized temperature sensitive septin mutant, *cdc12-6* (Adams and Pringle, 1984; Johnson *et al*., 2015) (Fig.5). In these mutants, septins localize normally to the bud-neck at permissive temperature, but rapidly disassemble at restrictive temperature (Gladfelter *et al*., 2005). First, we evaluated whether the AH was necessary for curvature sensing using recombinant septin complex with the cdc12-6 protein. Interestingly, we found that cdc12-6 mutant complexes bound all tested membrane curvatures indiscriminately in contrast to wild-type septins (Fig.5C and D). This suggests that AH of Cdc12 is not required for septin membrane association, but rather it specifies septin preference for membrane-curvature.

Next, we asked if the AH of Cdc12 was sufficient to distinguish membrane curvature. We purified a polypeptide with two copies of the AH to mimic the stoichiometry of the intact septin complex which has two copies of Cdc12. This tandem AH polypeptide adsorbed onto membrane curvatures of 2 μm^−1^ (albeit significantly less than septin complexes), while showing reduced binding to other tested membrane curvatures of 6.67 μm^−1^,4 μm^−1^, 0.67 μm^−1^ and 0.4 μm^−1^ (Fig.5E). The reduced binding onto beads of κ= 0.67 μm^−1^ was surprising, given the robust binding of septin complexes to this curvature (Fig. 1). We suspect that the spacing between tandem AH domains is an important factor in tuning the curvature preference as the spacing between AH domains in this construct is only 9 amino acids, whereas the spacing within a septin complex is approximately 24 nm (Bertin *et al*., 2008). Interestingly, a single AH-GFP adsorbed onto a range of more highly curved surfaces (κ=6.67, 4, and 2 μm^−1^) but less so on shallower curvatures (κ = 0.67 and 0.4 μm^−1^) (Supplemental Fig.2). However, we observed filament-like structures forming within both 2x AH-GFP and 1x AH-GFP constructs (Fig.5E and Supplemental Fig.2K, white arrows), suggesting that AH domains are capable of oligomerizing at the membrane, which might be expected given the small size of the domains (Huang and Ramamurthi, 2010). We see identical behavior with a tandem construct lacking a GFP tag and visualized by dye labeling showing that the preference and any oligomerization is not an artifact of the tag (unpublished result). Collectively, this data suggests that the spatial arrangement of AH domains and the capacity to self-assemble likely tune curvature sensitivity.

There is strong evidence that AH domains demonstrate lipid-specific binding (Vanni *et al*., 2014; Hofbauer *et al*., 2018). Similarly, septins preferentially bind membranes containing phosphatidylinositol (4,5) bis-phosphate (PI_4,5_P_2_), which is dependent on the presence of polybasic regions within individual septin polypeptides (Casamayor and Snyder, 2003; Akil *et al*., 2016). We probed the AH domain lipid-specificity by measuring adsorption onto both optimum (κ = 2 μm^−1^) and non-optimum (κ = 0.4 μm^−1^) membrane curvatures with varying lipid compositions while keeping the global charge across the membrane equivalent. There are, local differences in charge of the lipids used with PI_4,5_P_2_ (−3 at pH 7.4) greater than PI or PS (both −1 at pH 7.4) (Tsui, Ojcius and Hubbell, 1986; Beber *et al*., 2018). We found that curvature sensitivity remained intact on the different lipids, however, we observed increased binding of the 2x-AH domain, the single AH-GFP and the full septin complex to PI_4,5_P_2_containing membranes over membranes containing either phosphatidylinositol (PI) or phosphotidylserine (PS) (Fig. 5F, Supplemental Fig.2K,L,M and N). Interestingly, we observed stronger binding of the 2x-AH to PI over PS, despite their equal charge, suggesting that either the molecular shape of the lipid or the different fatty acyl chain environment influences AH adsorption.

Finally, we assessed the functionality of the Cdc12 AH in live cells using *Ashbya gossypii*, a filamentous fungus which displays prominent septin assemblies at sites of micron-scale curvature (DeMay *et al*., 2009; Bridges *et al*., 2016). We generated a *cdc12-6* mutant allele in *Ashbya*, similar to the *S. cerevisiae* allele. In *cdc12-6* mutants, septins rarely assemble at branches (17%; n=104 branches;12 cells), but still assembled into tightly-bundled filaments, septation site rings and thin filaments at hyphal tips (Fig. 5G). The few branch assemblies detected were aberrant, with low signal intensity and only at the base of newly formed branches. This shows that the amphipathic helix in Cdc12 is required for septin localization at curved membranes in cells.

### Conclusion

We identify a mechanism that uses both affinity and the number of available binding sites to drive cooperative, curvature-specific accumulation of septins. Even a single septin octamer is capable of detecting changes micrometer scale curvatures, manifested as differences in association rate. An amphipathic helix located at the very C-terminus of Cdc12 is necessary and sufficient for septins to distinguish different lipid compositions and curvatures. It is now clear that AH domains are not just useful for sensing nanometer-scale curvature but also to detect micrometer-scale curvature. The ability of septins to bind membranes in the absence of the AH domain suggests that there are additional curvature-independent septin-membrane interaction motifs. What septin amphipathic helices sense in the local lipid environment will be an exciting area of future investigation.

## Materials and Methods

### Yeast septin purification

BL21 (DE3) *E. coli* cells were transformed using a duet expression system (*Bridges et al*. 2016) and selected with ampicillin and chloramphenicol. Selected cells were cultured to an O.D._600 nm_ between 0.6 and 0.8 and induced with 1 mM of IPTG. Induced cultures were grown for 24 hours at 22° C (3 hours for Cdc12 amphipathic helix construct) before harvesting. Cultures were pelleted at 10,000 RCF for 15 min. Pellets were resuspended in lysis buffer (1M KCl, 50 mM, Hepes pH 7.4, 1 mM MgCl_2_, 10% glycerol, 1% Tween-20, and 1x protease inhibitor tablet (Roche), 20 mM Imidazole, 1 mg/mL lysozyme) for 30 minutes on ice with intermittent vortexing. The lysate was sonicated twice for 10 seconds and clarified by centrifugation using an SS-34 rotor at 20,000 RPM for 30 minutes. Clarified supernatant was filtered using 0.44 μm filter, then incubated with equilibrated Nickel-NTA^2+^ or Cobalt resin (Thermo Fisher Scientific; 2 mL resin per liter of *E*. coli culture) at 4°C for 1 hour. Resin and lysate were added to a gravity flow column. Bound protein was washed four times with 5x the column volume with wash buffer (1M KCl, 50 mM Hepes pH 7.4, 20 mM Imidazole), and then eluted in elution buffer (300mM KCl, 50 mM Hepes pH 7.4, 500 mM Imidazole). Eluted protein was then dialyzed into septin storage buffer (300 mM KCl, 50 mM Hepes pH 7.4, 1 mM BME) for 24 hours in two steps (Slide-A-Lyzer G2 20K MWCO, Thermo Fisher Scientific; 10K MWCO for amphipathic helix constructs). During dialysis, 60 μg of TEV protease (Sigma) was added to cleave the 6x-histidine tag on Cdc12. 24 hours later, the protein was run over a second column (either Ni-NTA or cobalt resin) to remove the protease, the poly-histidine tag, and additional contaminants. Protein purity was determined via SDS-PAGE and protein concentration was determined via Bradford assay. Purification of the Cdc12-6 complexes and AH domain constructs were purified as described above. However, for AH domain purifications, cells were induced for 4 hours at 37° C and harvested for protein purification.

### Amphipathic helix construct sequences

2x-AH-GFP construct has the following sequence followed by a GFP tag:

<u>LEEIQGKVKKLEEQVKSL</u>-GSGSRSGSGS-<u>LEEIQGKVKKLEEQVKSL</u>-GSGSSR-GFP tag, where the underlined sequences are the AH domains within Cdc12 and the serine-glycine repeats constitute the linker regions between tandem AH domains and the GFP tag. Similarly, the 1x-AH construct has the following sequence: <u>LEEIQGKVKKLEEQVKSL</u>-GSGSSR-GFP tag.

### Lipid mix preparation

All septin binding experiments were done using supported lipid bilayers consisting of 75 mol % dioleoylphosphatidylcholine (DOPC) Avanti Polar Lipids, 25 mol % PI (Liver, Bovine) sodium salt, and trace amounts (less than 0.1%) of phosphatidylethanolamine-N-(lissamine rhodamine B sulfonyl) (Rh-PE) (ammonium salt) Egg-Transphosphatidylated, Chicken) (Avanti Polar Lipids 810146) unless otherwise mentioned. All lipid compositions were made the same way. Lipids were mixed in chloroform solvent, and dried by a stream of argon gas to generate a lipid film. Remaining solvent was evaporated by placing the lipid film under a vacuum for at least 6 hours. Lipids were rehydrated in supported lipid bilayer buffer (300 mM KCl, 20 mM Hepes pH 7.4, 1 mM MgCl_2_) for 30 minutes at 37°C to give a final concentration of 5 mM. Hydrated lipids were subject to vortex for 10 seconds, every 5 mins for 30 mins bath sonicated in 2 minute intervals until clarification to yield small unilamellar vesicles (SUVs).

### Preparation of supported lipid bilayers on silica microspheres

SUVs were adsorbed onto silica microspheres (Bangs laboratories) of various curvatures by mixing 50 nM lipids with 440 um^2^ of silica microsphere surface area or 10 uL of rods in a final volume of 80 uL for 1 hour on a roller drum at room temperature. Unbound lipid was washed away by pelleting lipid-coated beads at the minimum force required to pellet each bead size using pre-reaction buffer (33.3 mM KCl, 50 mM Hepes pH 7.4) (See http://www.bangslabs.com/ for sedimentation speeds). Washes were preformed 4 times.

### Measuring protein adsorption onto silica microspheres

To examine septin adsorption onto different membrane curvatures, 25 μL of septins in septin storage buffer were added to 75 μL of a bead-buffer solution of 5 mm^2^ total lipid-bead surface area to give a final buffer composition of 100 mM KCl, 50 mM Hepes pH 7.4, 1 mM BME, 0.1% methylcellulose, and 0.1% bovine serum albumin (fatty acid free-Sigma). Saturation binding curves were obtained using several different septin concentrations in simple mixtures, containing only one bead size per reaction volume. Amphipathic helix adsorption was measured in mixtures containing protein and various bead diameters. Each reaction was incubated in a plastic chamber (Bridges methods paper) glued to a polyethyleneglycol (PEG)-passivated coverslip for 1 hour (to allow the reaction to reach equilibrium). Beads were imaged using either a wide-field microscope with a Ti-82 Nikon stage a 100x Plan Apo 1.49 NA oil lens, and Zyla sCMOS (Andor) (saturation binding curves) or a spinning disc (Yokogawa) confocal microscope (Nikon Ti-82 stage) using a 100x Plan Apo 1.49 NA oil lens, and a Prime 95B CMOS (Photometrics) (amphipathic helix and Cdc12-6 septin complex adsorption experiments). For analysis of septin binding, raw images were exported into Imaris 8.1.2 (Bitplane AG, Zurich, Switzerland). Each image was background subtracted in both channels using the software’s Gaussian filter for background determination (width 31.4 μm). The surface of each bead was defined using the lipid channel. Using the surface generate on each bead, sum intensity values from lipid and septin channels were exported from Imaris into Microsoft Excel. Septin adsorption was calculated by dividing the multiple sum intensity values from the septin channel over the sum intensity values from the lipid channel as to control for surface area for each bead size. Boxplots were generated through exporting intensity sum values from both channels into R version 3.2. 2 (R Foundation for Statistical Computing, Vienna, Austria, using RStudio 0.99.467, Boston, MA, USA). Boxplots were generated using ggplot2 package (Wickham, 2007; Wickham, 2009; Winston, 2014).

### Generation and preparation of septin-rod supported lipid bilayer mixture

Borosilicate rods were obtained from Glass Microfiber filters, GFC, 42.5 mm (Whatman). A single filter was torn up into small pieces into a beaker with 60 mL of 100% ethanol and sonicated until the solution became opaque. The solution was stored at room temperature overnight. The next day, 10 μL of rods were taken from the solution after thorough mixing. 70 uL of SLB buffer was added to the rods and spun down at top speed to dilute the ethanol in the solution. This step was repeated 4 additional times. 5 mM of SUVs (75% DOPC, 25% PI, trace Rh-PE, as described above) were added to the polycarbonate rods and allowed to incubate for 1 hour at room temperature. Unbound lipid was washed away pelleting lipid-coated beads at top speed (16.1 RCF) using pre-reaction buffer. The mixture of septin-lipid-coated rods were added to a circular 12mm (PEG)-coated coverslips and incubated at room temperature for 1 hour and then and prepared for scanning electron microscopy.

### Scanning electron microscopy

The septin-rod mixture onto a circular PEG-coated 12mm coverslip, was fixed in 2.5% glutaraldehyde in 0.05 M sodium cacodylate (NaCo) pH 7.4 for 30 minutes followed by 2x washes in 0.05 M NaCo (5 min each wash). Samples were post-fixed in 0.5% OsO_4_ Cacodylate buffer for 30 minutes and washed 3x in NaCo (5 minutes each wash). Samples were then incubated with 1% tannic acid for 15 minutes followed by 3x washes in NaCo. 0.5% OsO_4_ was added for 15 minutes followed by 3x washes in NaCo. Samples were then dehydrated with increasing ethanol concentrations: (30% EtOH for 5 minuets, 2x; 50% EtOH for 5 minutes; 75% EtOH for 5 minutes; 100% EtOH for 5 minuetes, 2x followed by another 10 minute incubation). Samples were incubated in transition fluid (hexamethyldisilazane) 3x; (incubation times: 5 minutes, 10 minutes, 5 minutes) and were allowed to airdry and then placed in a desiccator until sputter coating. Samples were coated in a gold/palladium alloy and then imaged on a Zeiss Supra 25 Field Emission Scanning Electron Microscope.

### Kinetics of single septin complex onto lipid-coated beads

Two PEG-passivated coverslips were sandwiched together using double-coated pressure sensitive adhesive tape (Nitto product:5015ELE) to make narrow (~20 μL flow chambers). A mixture of septins and lipid-coated beads of a given diameter were then flowed through the chamber and imaged using near-total internal reflection fluorescence microscopy. The number and duration of binding events were performed manually. Association-rate was quantified by calculating the number of binding events over the product of septin concentration, time, and binding area. To ensure the off rates were not dominated by bleaching, we generated bleach profiles (Supplemental fig. 1). The amount of time it took to bleach single septin complexes is much higher than the observed dwell time (~ 1.5 seconds). Bleach steps were also used to calibrate intensity so that scored puncta were single septin complexes. As a control, we measured the dwell time of single septin molecules that localized to areas of the coverslip that did not contain any beads and measured a dwell time of 0.062 seconds (data not shown), ensuring that our measured dwell times on beads reflect accurate association dynamics of single septin molecules on lipid-coated beads.

### Generation of helical wheel diagrams

Helical wheels and the calculation of net charge and hydrophobic moment were generated using Heliquest (http://heliquest.ipmc.cnrs.fr).

### *Ashbya gossypii* strain construction, culture and imaging

We generated an analogous *cdc12-6* allele (LEEVQAKVKKLEEQVRALQLRKH* ➔ LEEVQAKVKN*) with GFP tag for integration at the endogenous locus by cloning a FragmentGENE (Genewiz) (637bp) harboring the mutation into AGB260 (*pCDC12*-GFP:*GEN*) (Meseroll, Occhipinti and Gladfelter, 2013) via the restriction enzymes *BspEI* and *BsrGI* yielding AGB1209 (*pcdc12-6*-GFP:*GEN*). AGB1209 was linearized by digestion with *SphI* for targeted integration at the *CDC12* locus, and transformed into the wild-type *A. gossypii* strain (Altmann-Jöhl and Philippsen, 1996) to generate AG884 (*Agcdc12-6*-GFP:GEN *leu2*Δ *thr4*Δ). AG384 (*AgCDC11a*-GFP:*GEN leu2*Δ *thr4*Δ) is described in (Meseroll, Occhipinti and Gladfelter, 2013).

*A. gossypii* strains were grown from spores in full medium at either 24°C (*cdc12-6*) for 24 h or 30°C (wt) for 16 h before harvesting mycelial cells. Cells were mounted onto Low Fluorescence medium pads solidified with 2% agarose for imaging. Images were acquired using a spinning disc (Yokogawa) confocal microscope (Nikon Ti-82 stage) using a 100x Plan Apo 1.49 NA oil lens, and a Prime 95B CMOS (Photometrics).

## Acknowledgements

We thank the Gladfelter lab and Danny Lew for useful discussions, Matthias Garten for ideas in setting up the rod assay, the UNC EM facility (Victoria Madden and Kristen White) for support with SEM and HHMI Faculty Scholars award to ASG.

## Author contributions

KSC, BLW conducted experiments, BLW and JMC constructed strains for experiments, BLW and JMC constructed plasmids for experiments, KSC, BLW, and ASG designed experiments.

## Abbreviations

SLB: Supported lipid bilayer
PEG: Poly-ethylene glycol
DOPC: Dioleoyl-phosphocholine
DOPS: Dioleoyl-phosphoserine
DOPE: Dioleoyl-phosphoethananolamine
PI: Phosphatidylinositol
PI(_4,5_)P_2_: Phosphatidylinositol-4,5-bis-phosphate
Rh-PE: Rhodamine phosphoethanolamine

## Supplemental Figure legends

**S.1. Single molecule photobleaching profiles and protein purification**

(A) Two-step photobleaching profile for a single septin complex. (B-D). Timelapse imaging of single septin molecules binding to a 1 μm bead (B). A single septin complex is bound to the bead. (C). A second septin complex binds to the bead, adjacent to the initially bound septin particle. (D). A single septin particle disassociates from the bead. Signal intensity profiles for (B-D) were all fit to Gaussian distributions. (E-G) Coomassie staining of recombinantly expressed 2x Cdc12-AH-GFP, 1x Cdc12 AH-GFP, and yeast septins Cdc11-GFP, Cdc3, Cdc12-6, Cdc10 after cleavage of 6xHIS tag via TEV-protease, respectively.

**S.2. The effects of lipid composition on septin filament orientation and binding of 1x-AH domain and septin complexes onto curved membranes.**

Supported lipid bilayers of various lipid compositions were generated onto borosilicate rods. 50 nM of purified septins were added to supported lipid bilayers and images were acquired using a scanning electron microscope. Representative images were selected from three different categories (100-400 nm, 401-600 nm, and 601-2300 nm. A subset of filaments were false colored; pink filaments are more parallel to the long axis of curvature; yellow filaments are oriented at ~ 45°; blue filaments are aligned to the axis of principal curvature (A-C). Representative images of septin filament alignment on DOPC (75%) and DOPS (25%)-coated rods. Rod diameters are 195 nm, 470 nm, and 1096 nm, respectively. (D-F). Representative images of septin filament alignment on DOPC (75%), PI (20%), and PI(4,5)P2 (5%)-coated rods. Rod diameters are 95 nm, 494 nm, and 1006 nm, respectively. (G-I). Representative images of septin filament alignment on DOPC (70%), DOPS (10%), DOPE (10%), and PI(4,5)P2 (10%)- coated rods. Rod diameters are 189 nm, 519 nm, and 1803 nm, respectively. Scale bars all 100 nm. (J) Box and whisker plot quantifying adsorption of 1 μM 1x-AH-GFP onto various membrane curvatures with a lipid composition of 75% DOPC, 25% PI, and trace Rh-PE. n > 74 beads per curvature. Black bars represent the median and error bars represent the standard deviation (K) Binding of 1 μM 1x-AH-GFP (green) to either κ =2μm^−1^ or 0.4μm^−1^ membrane curvatures with various lipid compositions. All images are contrasted identically. White arrows highlight filament-like structures. Scale bar 5 μm. (L) Box and whisker plot quantifying adsorption of 1x-AH-GFP onto membrane curvatures with various lipid compositions from (K). n > 74 beads per condition. Black bars represent the median and error bars represent the standard deviation (M) Adsorption of 15 nM of septin complex (green) to either κ =2μm^−1^ or 0.4μm^−1^ membrane curvatures with various lipid compositions. All images are contrasted identically. Scale bar 5 μm. (N) Box and whisker plot quantifying adsorption of septin complexes onto membrane curvatures with various lipid compositions from (M). n > 36 beads per condition Black bars represent the median and error bars represent the standard deviation.

**S.3. Septin amphipathic helices net charge and hydrophobicity**

(A-B) Net charge and hydrophobic moment of selected septin amphipathic helices, respectively. (C) Primary sequence alignment of septin amphipathic helices. (D) Helical wheel diagrams for selected membrane binding and/ or curvature sensing amphipathic helices. (E-F) Net charge and hydrophobic moment of selected amphipathic helices. Interestingly, AH domains with a net charge of zero (*S. cerevisiae, C. albicans, A. gossypii*, and *H. sapien*) have higher hydrophobic moments than AH domains that are charged (*C. elegans* and *D. melanogaster*). The properties of the AH domain might be tuned to reflect differences in lipid compositions across species, yet still localize to similar membrane geometries. The septin AH hydrophobic moments closely match SpoVM and DivIVA, other micron-scale sensors from bacteria.

## Literature Cited

Adams, A. E. M. and Pringle, J. R. (1984) ‘Relationship of actin and tubulin distribution to bud growth in wild-type and morphogenetic-mutant Saccharomyces cerevisiae’, Journal of Cell Biology. doi: 10.1083/jcb.98.3.934.

Akil, A. et al. (2016) ‘Septin 9 induces lipid droplets growth by a phosphatidylinositol-5-phosphate and microtubule-dependent mechanism hijacked by HCV’, Nature Communications. doi: 10.1016/0021-8502(80)90042-7.

Altmann-Jöhl, R. and Philippsen, P. (1996) ‘AgTHR4, a new selection marker for transformation of the filamentous fungus Ashbya gossypii, maps in a four-gene cluster that is conserved between A. gossypii and Saccharomyces cerevisiae’, Molecular and General Genetics. doi: 10.1007/BF02191826.

Beber, A. et al. (2018) ‘Septin-based readout of PI(4,5)P2 incorporation into membranes of giant unilamellar vesicles’, Cytoskeleton. doi: 10.1002/cm.21480.

Bertin, A. et al. (2008) ‘Saccharomyces cerevisiae septins: Supramolecular organization of heterooligomers and the mechanism of filament assembly’, Proceedings of the National Academy of Sciences. doi: 10.1073/pnas.0803330105.

Bridges, A. A. et al. (2016) ‘Micron-scale plasma membrane curvature is recognized by the septin cytoskeleton’, Journal of Cell Biology. doi: 10.1083/jcb.201512029.

Byers, B. and Goetsch, L. E. (1976) ‘A highly ordered ring of membrane-associated filaments in budding yeast’, Journal of Cell Biology. doi: 10.1083/jcb.69.3.717.

Cannon, K. S., Woods, B. L. and Gladfelter, A. S. (2017) ‘The Unsolved Problem of How Cells Sense Micron-Scale Curvature’, Trends in Biochemical Sciences. doi: 10.1016/j.tibs.2017.10.001.

Casamayor, A. and Snyder, M. (2003) ‘Molecular dissection of a yeast septin: distinct domains are required for septin interaction, localization, and function.’, Molecular and cellular biology. doi: 10.1128/MCB.23.8.2762.

Cho, S. J. et al. (2011) ‘Septin 6 regulates the cytoarchitecture of neurons through localization at dendritic branch points and bases of protrusions’, Molecules and Cells. doi: 10.1007/s10059-011-1048-9.

Clay, L. et al. (2014) ‘A sphingolipid-dependent diffusion barrier confines ER stress to the yeast mother cell’, eLife. doi: 10.7554/eLife.01883.

DeMay, B. S. et al. (2009) ‘Regulation of distinct septin rings in a single cell by Elm1p and Gin4p kinases.’, Molecular biology of the cell. doi: 10.1091/mbc.E08-12-1169.

Drin, G. et al. (2007) ‘A general amphipathic α-helical motif for sensing membrane curvature’, Nature Structural and Molecular Biology. doi: 10.1038/nsmb1194.

Drin, G. and Antonny, B. (2010) ‘Amphipathic helices and membrane curvature’, FEBS Letters. doi: 10.1016/j.febslet.2009.10.022.

Field, C. M. et al. (1996) ‘A purified Drosophila septin complex forms filaments and exhibits GTPase activity’, Journal of Cell Biology. doi: 10.1083/jcb.133.3.605.

Finnigan, G. C. et al. (2015) ‘The carboxy-terminal tails of septins Cdc11 and Shs1 recruit myosin-II binding factor Bni5 to the bud neck in Saccharomyces cerevisiae’, Genetics. doi: 10.1534/genetics.115.176503.

Ford, S. K. and Pringle, J. R. (1991) ‘Cellular morphogenesis in the Saccharomyces cerevisiae cell cycle: Localization of the CDC11 gene product and the timing of events at the budding site’, Developmental Genetics. doi: 10.1002/dvg.1020120405.

Frazier, J. A. et al. (1998) ‘Polymerization of purified yeast septins: Evidence that organized filament arrays may not be required for septin function’, Journal of Cell Biology. doi: 10.1083/jcb.143.3.737.

Garcia, G. et al. (2011) ‘Subunit-dependent modulation of septin assembly: Budding yeast septin Shs1 promotes ring and gauze formation’, Journal of Cell Biology. doi: 10.1083/jcb.201107123.

Gautier, R. et al. (2008) ‘HELIQUEST: A web server to screen sequences with specific α-helical properties’, Bioinformatics. doi: 10.1093/bioinformatics/btn392.

Gill, R. L. et al. (2015) ‘Structural basis for the geometry-driven localization of a small protein’, Proceedings of the National Academy of Sciences. doi: 10.1073/pnas.1423868112.

Gladfelter, A. S. et al. (2005) ‘Interplay between septin organization, cell cycle and cell shape in yeast.’, Journal of cell science. doi: 10.1242/jcs.02286.

Gopalakrishnan, G. et al. (2009) ‘Supported bilayers formed from different phospholipids on spherical silica substrates’, Langmuir. doi: 10.1021/la9006982.

Haarer, B. K. and Pringle, J. R. (1987) ‘Immunofluorescence localization of the Saccharomyces cerevisiae CDC12 gene product to the vicinity of the 10-nm filaments in the mother-bud neck.’, Molecular and cellular biology. doi: 10.1128/mcb.7.10.3678.

Hatzakis, N. S. et al. (2009) ‘How curved membranes recruit amphipathic helices and protein anchoring motifs’, Nature Chemical Biology. doi: 10.1038/nchembio.213.

Hofbauer, H. F. et al. (2018) ‘The molecular recognition of phosphatidic acid by an amphipathic helix in Opi1.’, The Journal of cell biology. doi: 10.1083/jcb.201802027.

Huang, K. C. and Ramamurthi, K. S. (2010) ‘Macromolecules that prefer their membranes curvy.’, Mol Microbiol. doi: 10.1111/j.1365-2958.2010.07168.x.

Hussain, S. et al. (2018) ‘MreB filaments align along greatest principal membrane curvature to orient cell wall synthesis’, eLife. doi: 10.7554/eLife.32471.

John, C. M. et al. (2007) ‘The Caenorhabditis elegans septin complex is nonpolar’, EMBO Journal. doi: 10.1038/sj.emboj.7601775.

Johnson, C. R. et al. (2015) ‘Cytosolic chaperones mediate quality control of higher-order septin assembly in budding yeast’, Molecular Biology of the Cell. doi: 10.1091/mbc.E14-11-1531.

Joo, E., Surka, M. C. and Trimble, W. S. (2007) ‘Mammalian SEPT2 Is Required for Scaffolding Nonmuscle Myosin II and Its Kinases’, Developmental Cell. doi: 10.1016/j.devcel.2007.09.001.

Khan, A., Newby, J. and Gladfelter, A. S. (2018) ‘Control of septin filament flexibility and bundling by subunit composition and nucleotide interactions’, Molecular Biology of the Cell. doi: 10.1091/mbc.E17-10-0608.

Kim, E. Y. et al. (2017) ‘Dash-and-Recruit Mechanism Drives Membrane Curvature Recognition by the Small Bacterial Protein SpoVM’, Cell Systems. doi: 10.1016/j.cels.2017.10.004.

Longtine, M. S. et al. (2000) ‘Septin-Dependent Assembly of a Cell Cycle-Regulatory Module in Saccharomyces cerevisiae’, Molecular and Cellular Biology. doi: 10.1128/MCB.20.11.4049-4061.2000.

Maddox, A. S. et al. (2007) ‘Anillin and the Septins Promote Asymmetric Ingression of the Cytokinetic Furrow’, Developmental Cell. doi: 10.1016/j.devcel.2007.02.018.

Meitinger, F. et al. (2011) ‘Phosphorylation-dependent regulation of the F-BAR protein Hof1 during cytokinesis’, Genes and Development. doi: 10.1101/gad.622411.

Meseroll, R. A., Occhipinti, P. and Gladfelter, A. S. (2013) ‘Septin phosphorylation and coiled-coil domains function in cell and septin ring morphology in the filamentous fungus Ashbya gossypii’, Eukaryotic Cell. doi: 10.1128/EC.00251-12.

Pan, F., Malmberg, R. L. and Momany, M. (2007) ‘Analysis of septins across kingdoms reveals orthology and new motifs’, BMC Evolutionary Biology. doi: 10.1186/1471-2148-7-103.

Pranke, I. M. et al. (2011) ‘α-Synuclein and ALPS motifs are membrane curvature sensors whose contrasting chemistry mediates selective vesicle binding’, Journal of Cell Biology. doi: 10.1083/jcb.201011118.

Ramamurthi, K. S. et al. (2009) ‘Geometric cue for protein localization in a bacterium’, Science. doi: 10.1126/science.1169218.

Sadian, Y. et al. (2013) ‘The role of Cdc42 and Gic1 in the regulation of septin filament formation and dissociation’, eLife. doi: 10.7554/eLife.01085.

Sakchaisri, K. et al. (2004) ‘Coupling morphogenesis to mitotic entry.’, Proceedings of the National Academy of Sciences of the United States of America. doi: 10.1073/pnas.0400641101.

Simunovic, M. et al. (2015) ‘When Physics Takes Over: BAR Proteins and Membrane Curvature’, Trends in Cell Biology. doi: 10.1016/j.tcb.2015.09.005.

Simunovic, M., Srivastava, A. and Voth, G. A. (2013) ‘Linear aggregation of proteins on the membrane as a prelude to membrane remodeling’, Proceedings of the National Academy of Sciences. doi: 10.1073/pnas.1309819110.

Sirajuddin, M. et al. (2007) ‘Structural insight into filament formation by mammalian septins’, Nature. doi: 10.1038/nature06052.

Spiliotis, E. T., Kinoshita, M. and Nelson, W. J. (2005) ‘A mitotic septin scaffold required for mammalian chromosome congression and segregation’, Science. doi: 10.1126/science.1106823.

Tsui, F. C., Ojcius, D. M. and Hubbell, W. L. (1986) ‘The intrinsic pKa values for phosphatidylserine and phosphatidylethanolamine in phosphatidylcholine host bilayers’, Biophysical Journal. doi: 10.1016/S0006-3495(86)83655-4.

Ursell, T. S. et al. (2014) ‘Rod-like bacterial shape is maintained by feedback between cell curvature and cytoskeletal localization’, Proceedings of the National Academy of Sciences. doi: 10.1073/pnas.1317174111.

Vanni, S. et al. (2014) ‘A sub-nanometre view of how membrane curvature and composition modulate lipid packing and protein recruitment’, Nature Communications. Nature Publishing Group, a division of Macmillan Publishers Limited. All Rights Reserved., 5, p. 4916. Available at: https://doi.org/10.1038/ncomms5916.

Westfall, P. J. and Momany, M. (2002) ‘Aspergillus nidulans septin AspB plays pre- and postmitotic roles in septum, branch, and conidiophore development’, Molecular Biology of the Cell. doi: 10.1091/mbc.

Yamada, S. et al. (2016) ‘Septin Interferes with the Temperature-Dependent Domain Formation and Disappearance of Lipid Bilayer Membranes’, Langmuir. doi: 10.1021/acs.langmuir.6b03452.

Zimmerberg, J. and Kozlov, M. M. (2006) ‘How proteins produce cellular membrane curvature’, Nature Reviews Molecular Cell Biology. doi: 10.1038/nrm1784.

